# Potential cost and combinatorial effects of commensal gut microbes on survival of individual honey bees

**DOI:** 10.64898/2025.12.12.694077

**Authors:** Masato Sato, Ryo Miyazaki

## Abstract

The western honey bee, *Apis mellifera*, is attracting attention as an experimental model to investigate general interactions between hosts and gut microbiota because of their noteworthy simple and stable bacterial community. Here, we systematically explore the effect of five core bacteria, *Snodgrassella alvi, Bifidobacterium asteroides*, *Bombilactobacillus* Firm-4, *Lactobacillus* Firm-5, and *Gilliamella apicola*, on the longevity of individual honey bees. Generating gnotobiotic bees harboring all 32 different combinations of the core bacteria, we found that bacterial colonization generally reduces the host lifespan. This indicates that despite commensal gut microbes coevolved with the host, their presence is in principle costly for survival of individual bees. Regression analysis including higher-order interactions reveals that bacterial interactions have significant impacts on the host lifespan and show a qualitative trend: 2- and 4-way interactions exhibit positive, while 3- and 5-way interactions are negative. Numerical simulations based on a general Lotka-Volterra model demonstrate that the observed trend is reproduced when competitive interactions dominate in the bacterial community. This study combining empirical and theoretical approaches suggests the general costs of harboring gut microbiota and the stabilization of the bacterial community by higher-order interactions, which will advance our understanding of how commensal gut microbes and their interactions specifically influence host physiology.

**Importance:** Gut microbes are recognized for their profound effects on the host physiology. They provide numerous benefits to their hosts, whereas gut dysbiosis leads to various disease or mental impairment. Although understanding of such association with hosts is of pivotal importance, many animals have complex gut microbiome comprised of hundreds of bacterial species, most of which cannot be cultured in the laboratory. This is a big challenge to disentangle intimate interactions of commensal microbes with their hosts. Here, we exploit characteristics of honey bee gut microbiota, which consists of only five core bacteria, to generate gnotobiotic bees harboring defined bacterial communities with all 32 (*i.e*., 2^5^) possible combinations. This large-scale analysis uncovers not only individual impacts of the core species but also combinatorial effects generated by their interaction on the host lifespan. Finally, using simple mathematical models, we explain that the combinatorial effects are largely derived from competitions between gut microbes.

## Observation

Connections between gut microbiota and host health have attracted great attentions (1, 2). A thorough evaluation of the impact of gut microbiota on their hosts should encompass effects of not only individual bacteria but also interactions among bacteria. However, the complexity of gut microbiota in vertebrates, including humans, poses considerable challenges to achieving comprehensive evaluations. The western honey bee, *Apis mellifera*, has emerged as a valuable model system for studying gut microbiota with several advantages (3, 4): (i) its microbiota is simple with five core phylotypes (*Snodgrassella alvi, Gilliamella, Bifidobacterium, Bombilactobacillus* Firm-4, and *Lactobacillus* Firm-5) occupying more than 90% of the bacterial composition (5, 6), (ii) these core phylotypes are prevalent worldwide (7, 8), (iii) microbiota-depleted (MD) bees can be generated (9, 10), and (iv) the core bacterial species are culturable in the laboratory so that they can be used to generate gnotobiotic bees (11–13). While some benefits of the core bacteria for honey bees have been investigated (6, 14–17), comprehensive examination of the effects of bacterial interactions on bee physiology is scarce.

To evaluate exhaustive impacts of gut microbiota on the lifespan of honey bees, we conducted survival test of gnotobiotic bees harboring all 32 (*i.e*., 2^5^) possible combinations of the five core species, *S. alvi* (Sa), *Gilliamella apicola* (Ga), *Bifidobacterium asteroids* (Ba), *Bombilactobacillus* Firm-4 (L4), and *Lactobacillus* Firm-5 (L5) (Fig 1A). Gnotobiotic bees were prepared by feeding microbiota-depleted (MD) bees with the core species, and their colonization 10 days after treatment was verified by qPCR (Fig S1). As the population density of social insects could affect individual mortality (18), individual bees were solely reared in the cage to avoid effects of density variation during observation (see Supplementary Methods). Survival curves showed that host’s survival generally declined in the presence of core species, compared to MD bees (Fig 1B and Table S1). In monoinoculation groups, Sa caused a significant decrease of survival (Fig 1B, *p* = 0.001, Table S1). Since *S. alvi* is known to utilize host-derived nutritional resources (19), this result suggests that the resource allocation can be cost for honey bees. Other species also showed slight declines in lifespan, although not statistically significant. In 2-species inoculation groups, combinations of Sa with either Ba, L4, or L5 significantly reduced host lifespan (Fig 1B; *p* = 0.007, *p* = 0.03, and *p* = 0.002, respectively, Table S1). However, Sa with Ga showed no significant decrease of lifespan (*p* = 0.231, Table S1), suggesting that Ga counteracts the negative effect of Sa. Decline in survival tended to become larger with inoculations of more than 3 species, especially with significant reductions of lifespan by all 4- and 5-species inoculations (Fig 1B). There was no significant difference between 5-species and gut homogenate inoculations (Fig 1B; *p* = 0.229, Table S1), indicating that effects of minor microbial species and strain-level diversity included in the homogenate are negligible. Overall, core gut microbes tended to reduce the survival of honey bees, despite their consistent symbiotic relationship with the host. This implies that honey bees may receive reciprocal benefits from the core microbes, forming a tradeoff. Conceivable benefits by harboring gut microbiota could be extension of nutritional availability to various food resources in the fluctuating wild environment (20), or protection against pathogens by enhancing host immunity (21, 22). These benefits are most likely risk-hedging strategies for unpredictable environment, although these should be undetectable and costs would rather appear in our experimental condition where sufficient foods supplied and no pathogenic threat.

**Figure 1.**
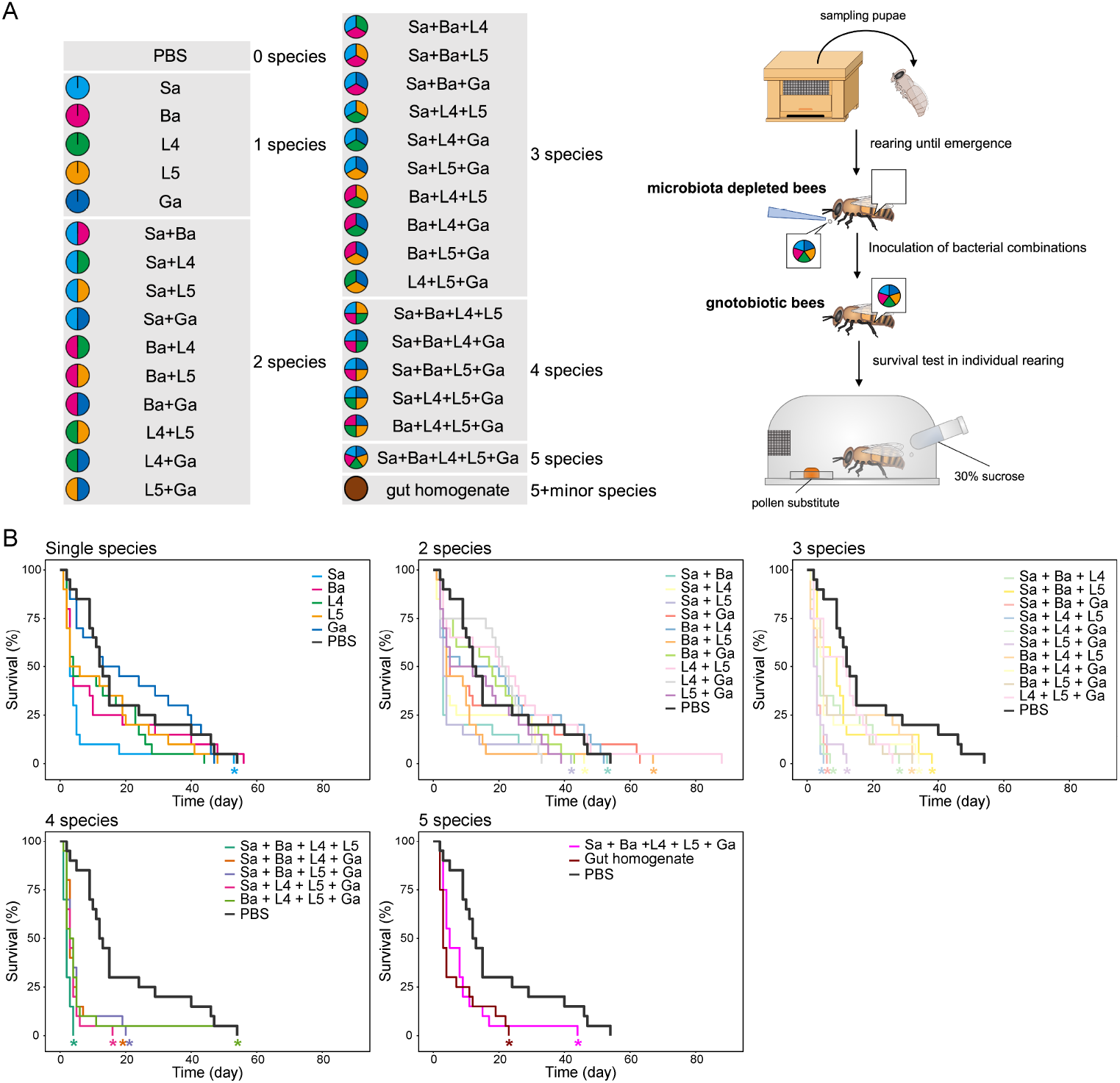
Survival of gnotobiotic honey bee workers with different combinations of core gut microbes. (A) Schematic illustrations of the experimental workflow. The list of all 32 different combinations of the the 5 core bacterial species, Sa (light blue), Ba (pink), L4 (green), L5 (yellow), and Ga (navy blue), along with the whole gut homogenate (brown), are shown on the left. The color of the pies indicates the containing species. A schematic illustration on the right shows the workflow for the survival test. The pupae were collected from hives and reared in the sterilized laboratory condition until they emerged, resulting in microbiota depleted (MD) bees. Bacterial cocktails were inoculated into the MD bees, and the gnotobiotic bees were reared individually to monitor their survival. A total of 20 bees from two colonies (10 bees from each) were tested for each treatment. (B) Inoculation of gut bacteria tends to decline the survival of honey bees. Each panels show Kaplan-Meier survival curves for inoculation treatments of single species, 2 species combinations, 3 species combinations, 4 species combinations, a 5 species combination, and a whole gut homogenate. All panels include the no-bacterial inoculation treatment (*i.e*., MD bees), indicated as PBS with a black bold curve. Statistical differences between different treatments are evaluated by generalized Wilcoxson log-rank test (Table S1), and significant differences (*p* < 0.05) compared to PBS treatment are indicated with an asterisk below each curve.

In principle, the outcomes of harboring multiple core microbes is the sum of the effects of individual microbes plus those arising from interactions among them. Our survival test with the all 32 combinations revealed the average lifespan of gnotobiotic bees (Fig 2A), allowing to estimate not only the impact of each core bacterium but also combinatorial effects emerging from specific bacterial combinations. Through GLMM regression analysis of average lifespan, we found that both effects of each bacterial species and higher-order combinations significantly influenced the host lifespan (Fig 2B and Table S2). In agreement with the survival curves (Fig 1B), all single-species effects were non-positive: Sa has the most negative impact on the lifespan (coefficient is −1.181, *p* < 0.001, Table S2), followed by L4, L5, and Ba (coefficients are −0.855, −0.752, and −0.669, *p* = 0.002, *p* = 0.008, and *p* = 0.013, respectively, Table S2), and only Ga has no significant effect (coefficient is −0.12, *p* = 0.64, Table S2). Combinatorial effects of multiple species exhibited a characteristic trend: 2- and 4-way interactions showed positive effects, while 3- and 5-way interactions had negative effects on the lifespan (Fig 2B). We hypothesize that this qualitative trend originates from interspecific interactions among the core bacteria, resulting in changes in their relative density in the gut. To examine which types of bacterial interactions reproduce the observed trend, we analyzed three models with different types (see Supplementary Methods). These models involves randomly sampled interspecific interaction coefficients, *α*_*ij*_, from various distributions: the first assumes all *α*_*ij*_ are negative (competition model); the second assumes all *α*_*ij*_ are positive (cooperation model); the third allows the sign of *α*_*ij*_can be positive or negative (random model). Numerical analysis revealed that the empirically observed trend was reproduced only in the competition model (Fig 2C). When two competitive species coexist, their densities must decrease compared to monocultures. Then, the effect of individual species on the host weakens as their densities decline. Since all individual species negatively (or non-positively) impact the host lifespan, the weakening of individual effects consequently leads to positive effects on the lifespan. Similarly, adding third species weakens the positive 2-way effects by causing further decline of densities of both species, resulting in negative 3-way effects.

**Figure 2.**
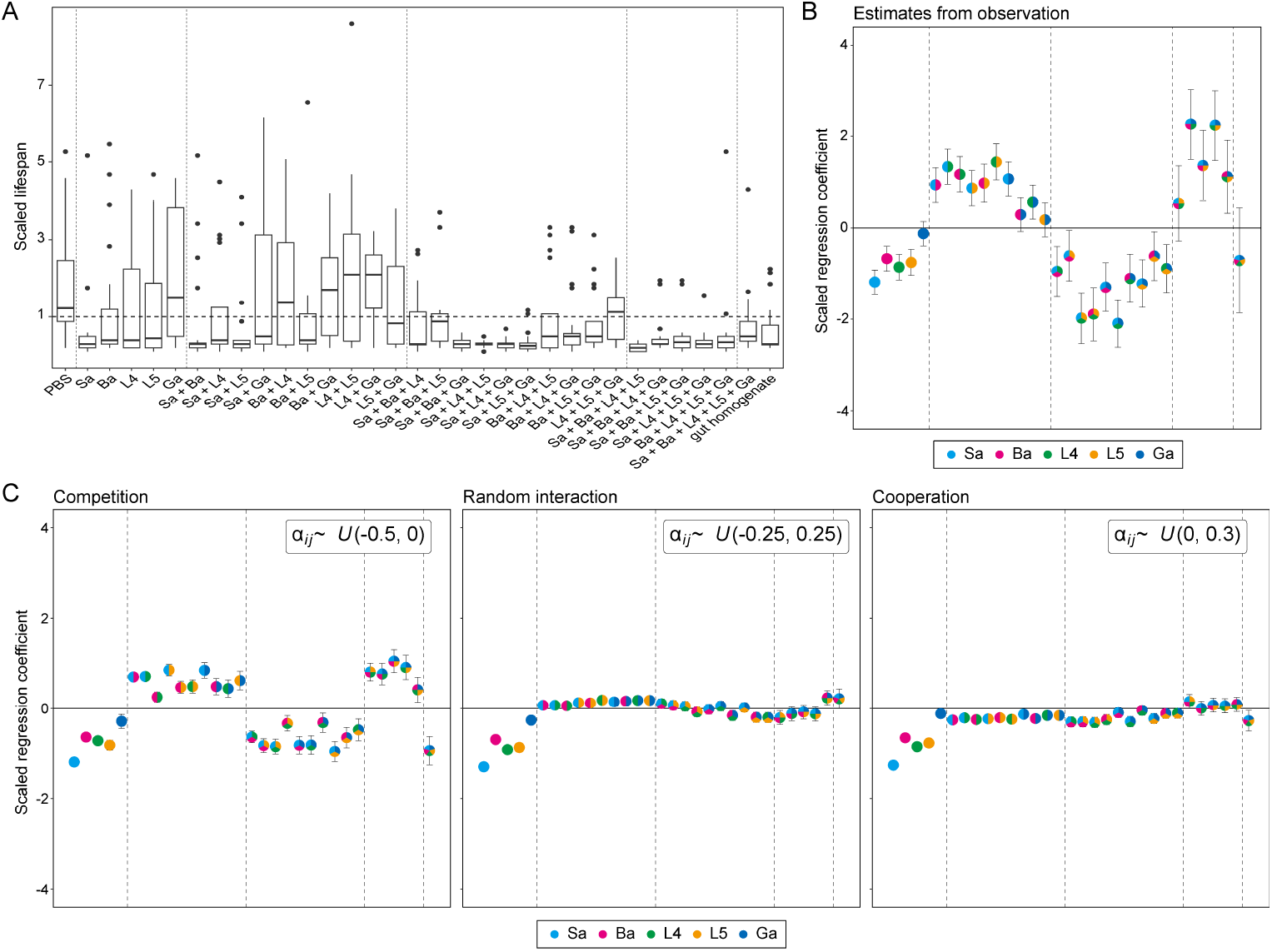
Net effects of interspecific interactions between core gut microbes on the bee lifespan. (A) Lifespans of gnotobiotic honey bee workers with different combinations of the core gut bacteria. Lifespans are scaled so that the mean of all samples is 1 (a horizontal broken line). Box plot indicates interquartile range with the bold line representing the median, and outliers more than 1.5 times out of the interquartile range. The vertical dotted lines separate the plots by the number of combination. (B) Scaled regression coefficients of each of bacteria and higher-order bacterial interactions. Regression coefficients are drowned by fitting a generalized linear mixed model using average lifespan as a response variable and inoculation of each of bacteria as the explanatory variable (indicator variable). Plotted values are scaled as the estimated intercept is 1. Colored pies indicate bacterial species and their higher-order interactions. Error bars represent the standard errors. Regression coefficients are scaled by the estimated value of intercept. Note that higher-order interactions coefficients indicate only the effect of interactions. The values of regression coefficients and the summary of GLMM analysis is shown in Table S2. (C) Regression coefficients estimated from simulations of various pairwise bacterial interactions. Regression coefficients are estimated from data generated by 1000 times numerical simulations of pairwise competition model (*α*_*ij*_∼*U*(−0.5, 0)) (left panel), pairwise random interactions model (*α*_*ij*_∼*U*(−0.25, 0.25)) (middle panel), and pairwise cooperation model (*α*_*ij*_∼*U*(0, 0.3)) (right panel). Colored pies indicate bacterial species and their higher-order interactions, as shown in B. Error bars represent the standard errors.

Previous studies on bacterial interactions in diverse communities including gut microbiota consistently highlights the prevalence of competition (23). Some members of honey bee gut microbiota show tremendous variability in the expression of type VI secretion systems associated toxins to kill neighboring bacterial cells (24), suggesting the importance of competitions within the gut. While competition among gut bacteria is plausible, empirical evidence also suggests cross-feeding between *S. alvi* and *G. apicola* (9, 15). In this study, interspecific interactions are considered as net effects, represented by the coefficient *α*_*ij*_ (see Supplementary Methods), and thus all individual interactions, including cross-feeding, are taken into account in the value. More detailed and explicit models of individual interactions will be developed when quantitative data are available on the resources used competitively or those exchanged through cross-feeding. Decline of host survival by harboring gut microbiota has been also reported in *Drosophila melanogaster* (25). Five gut bacterial species associated with fruit flies but not with honey bees were inoculated into *D. melanogaster* with all combinations, although they rather focused on the context-dependent effects of how pairwise or three-way interactions are modulated by the presence of bystander species (25). When we plotted regression coefficients using their data of the linear regression analysis, the similar qualitative trend to our current results of honey bees was observed (Fig S2). As demonstrated in our mathematical model, this trend could be reproduced when individual effects of the bacterial members on their hosts exhibit a consistent sign (*i.e*., either positive or negative), and competition is the dominant mode of interspecies interaction. Together with the ubiquity of competition among bacteria, our results would capture a fundamental aspect of bacterial interactions in commensal microbial communities.

## Acknowledgments

We thank Philipp Engel for providing bacterial strains under the collaborative work supported by Human Frontier Science Program (RGY0077/2016), and Wenying Shou for critical reading of the manuscript. We thank Keiko Takano, Kayo Ohkouchi, and Yuko Iijima for their technical assistance. This study was supported by Japan Science and Technology Agency FORESTO Program (JPMJFR201C) and Japan Society for the Promotion of Science (JSPS) KAKENHI (21H02213 and 24K01772). M.S. was supported by JSPS postdoctoral fellowship (202400875) and KAKENHI (24KJ2214).

## Author contributions

Conceptualization – RM; Methodology – MS, RM; Software – MS; Validation – MS; Formal analysis – MS; Investigation – MS; Resources – RM; Data curation – MS, RM; Writing - Original Draft – MS; Review and Editing – MS, RM; Visualization – MS, RM; Supervision – RM; Project administration – RM; Funding acquisition – MS, RM

## References

1. Grenham S, Clarke G, Cryan J, Dinan T. 2011. Brain–gut–microbe communication in health and disease. Front Physiol 2.

2. Shreiner AB, Kao JY, Young VB. 2015. The gut microbiome in health and in disease. Curr Opin Gastroenterol 31:69–75.

3. Engel P, James RR, Koga R, Kwong WK, McFrederick QS, Moran NA. 2013. Standard methods for research on Apis mellifera gut symbionts. J Apic Res 52:1–24.

4. Kwong WK, Moran NA. 2016. Gut microbial communities of social bees. Nat Rev Microbiol 14:374–384.

5. Cox-Foster DL, Conlan S, Holmes EC, Palacios G, Evans JD, Moran NA, Quan P-L, Briese T, Hornig M, Geiser DM, Martinson V, vanEngelsdorp D, Kalkstein AL, Drysdale A, Hui J, Zhai J, Cui L, Hutchison SK, Simons JF, Egholm M, Pettis JS, Lipkin WI. 2007. A metagenomic survey of microbes in honey bee colony collapse disorder. Science 318:283–287.

6. Martinson VG, Moy J, Moran NA. 2012. Establishment of characteristic gut bacteria during development of the honeybee worker. Appl Environ Microbiol 78:2830–2840.

7. Ahn J-H, Hong I-P, Bok J-I, Kim B-Y, Song J, Weon H-Y. 2012. Pyrosequencing analysis of the bacterial communities in the guts of honey bees Apis cerana and Apis mellifera in Korea. J Microbiol 50:735–745.

8. Sabree ZL, Hansen AK, Moran NA. 2012. Independent studies using deep sequencing resolve the same set of core bacterial species dominating gut communities of honey bees. PLOS ONE 7:e41250.

9. Kwong WK, Engel P, Koch H, Moran NA. 2014. Genomics and host specialization of honey bee and bumble bee gut symbionts. Proc Natl Acad Sci 111:11509–11514.

10. Engel P, Bartlett KD, Moran NA. 2015. The bacterium Frischella perrara causes scab formation in the gut of its honeybee host. mBio 6:e00193–15.

11. Scardovi V, Trovatelli LD. 1969. New species of bifid bacteria from Apis mellifica L. and Apis indica F. A contribution to the taxonomy and biochemistry of the genus Bifidobacterium. Zentralblatt Bakteriol Parasitenkd Infekt Hyg Zweite Naturwissenschaftliche Abt Allg Landwirtsch Tech Mikrobiol 123:64–88.

12. Kwong W, Moran N. 2012. Cultivation and characterization of the gut symbionts of honey bees and bumble bees: description of Snodgrassella alvi gen. nov., sp. nov., a member of the family Neisseriaceae of the Betaproteobacteria, and Gilliamella apicola gen. nov., sp. nov., a member of Orbaceae fam. nov., Orbales ord. nov., a sister taxon to the order “Enterobacteriales” of the Gammaproteobacteria. Int J Syst Evol Microbiol 63.

13. Olofsson TC, Alsterfjord M, Nilson B, Butler È, Vásquez A. 2014. Lactobacillus apinorum sp. nov., Lactobacillus mellifer sp. nov., Lactobacillus mellis sp. nov., Lactobacillus melliventris sp. nov., Lactobacillus kimbladii sp. nov., Lactobacillus helsingborgensis sp. nov. and Lactobacillus kullabergensis sp. nov., isolated from the honey stomach of the honeybee Apis mellifera. Int J Syst Evol Microbiol 64:3109–3119.

14. Horak RD, Leonard SP, Moran NA. 2020. Symbionts shape host innate immunity in honeybees. Proc R Soc B Biol Sci 287:20201184.

15. Kešnerová L, Mars RAT, Ellegaard KM, Troilo M, Sauer U, Engel P. 2017. Disentangling metabolic functions of bacteria in the honey bee gut. PLOS Biol 15:e2003467.

16. Ellegaard KM, Tamarit D, Javelind E, Olofsson TC, Andersson SG, Vásquez A. 2015. Extensive intra-phylotype diversity in lactobacilli and bifidobacteria from the honeybee gut. BMC Genomics 16:284.

17. Liberti J, Kay T, Quinn A, Kesner L, Frank ET, Cabirol A, Richardson TO, Engel P, Keller L. 2022. The gut microbiota affects the social network of honeybees. 10. Nat Ecol Evol 6:1471–1479.

18. Koto A, Mersch D, Hollis B, Keller L. 2015. Social isolation causes mortality by disrupting energy homeostasis in ants. Behav Ecol Sociobiol 69:583–591.

19. Quinn A, El Chazli Y, Escrig S, Daraspe J, Neuschwander N, McNally A, Genoud C, Meibom A, Engel P. 2024. Host-derived organic acids enable gut colonization of the honey bee symbiont Snodgrassella alvi. Nat Microbiol 9:477–489.

20. Engel P, Martinson VG, Moran NA. 2012. Functional diversity within the simple gut microbiota of the honey bee. Proc Natl Acad Sci 109:11002–11007.

21. Steele MI, Motta EVS, Gattu T, Martinez D, Moran NA. 2021. The gut microbiota protects bees from invasion by a bacterial pathogen. Microbiol Spectr 9:e00394–21.

22. Lang H, Duan H, Wang J, Zhang W, Guo J, Zhang X, Hu X, Zheng H. 2022. Specific strains of honeybee gut Lactobacillus stimulate host immune system to protect against pathogenic Hafnia alvei. Microbiol Spectr 10:e01896–21.

23. Palmer JD, Foster KR. 2022. Bacterial species rarely work together. Science 376:581–582.

24. Steele MI, Kwong WK, Whiteley M, Moran NA. 2017. Diversification of type VI secretion system toxins reveals ancient antagonism among bee gut microbes. mBio 8:e01630–17.

25. Gould AL, Zhang V, Lamberti L, Jones EW, Obadia B, Korasidis N, Gavryushkin A, Carlson JM, Beerenwinkel N, Ludington WB. 2018. Microbiome interactions shape host fitness. Proc Natl Acad Sci 115:E11951–E11960.

